# Φ-Space ST: a platform-agnostic method to identify cell states in spatial transcriptomics studies

**DOI:** 10.1101/2025.02.05.636735

**Authors:** Jiadong Mao, Jarny Choi, Kim-Anh Lê Cao

**Affiliations:** Melbourne Integrative Genomics, School of Mathematics and Statistics, The University of Melbourne, Australia; Bioinformatics and Cellular Genomics, St Vincent’s Institute, Australia

## Abstract

We introduce Φ-Space ST, a platform-agnostic method to identify continuous cell states in spatial transcriptomics (ST) data using multiple scRNA-seq references. For ST with supercellular resolution, Φ-Space ST achieves interpretable cell type deconvolution with significantly faster computation. For subcellular resolution, Φ-Space ST annotates cell states without cell segmentation, leading to highly insightful spatial niche identification. Φ-Space ST harmonises annotations derived from multiple scRNA-seq references, and provides interpretable characterisations of disease cell states by leveraging healthy references. We validate Φ-Space ST in three case studies involving CosMx, Visium and Stereo-seq platforms for various cancer tissues. Our method revealed niche-specific enriched cell types and distinct cell type co-presence patterns that distinguish tumour from non-tumour tissue regions. These findings highlight the potential of Φ-Space ST as a robust and scalable tool for ST data analysis for understanding complex tissues and pathologies.

## 1 Introduction

The rapid advancement of spatial transcriptomics (ST) technologies has transformed our ability to explore the spatial organisation of gene expression within tissues, offering unprecedented insights into cellular architecture and tissue microenvironments [1]. Technologies such as 10x Genomics Visium [2], NanoString CosMx [3] and BGI’s Stereo-seq [4] have allowed researchers to map gene expression at various spatial resolutions, ranging from subcellular to supercellular levels. These advances have unlocked opportunities to understand tissue organisation and cellular interactions in both healthy and disease states. However, achieving biologically meaningful annotations of continuous cell states remains a major hurdle, particularly across platforms with varying resolutions and technical specifications.

A common strategy for cell type annotation in ST data involves comparing expression profiles of query ST data to those in well-annotated scRNA-seq references [5, 6]. However, existing ST platforms lack exact single-cell resolution since they do not directly measure single cells [1, 7]. For supercellular resolution ST such as Visium [2], each measurement unit is a multi-cellular spot and hence may contain a mixture of cell identities. For subcellular resolution ST such as CosMx [3] and Stereo-seq [4], the minimal measurement unit is often much smaller than the size of a single cell. These subcellular measurement units can be aggregated into suitable sizes via simple spatial binning or sophisticated cell segmentation algorithms [8, 9]. However, the resulted bins or segmented cells may still contain fractions of transcripts from multiple neighbouring cells [9, 10].

To deal with mixing cell identities commonly seen in both supercellular and subcellular ST data, cell type deconvolution methods have been developed [11–16]. These deconvolution methods can estimate cell type abundances within each spatial measurement unit such as supercellular spots, segmented cells and spatial bins based on reference scRNA-seq datasets. In the remainder of this article, we will refer to these spatial measurements as *cell-like objects*. Existing cell type deconvolution methods have a high potential but face several limitations, including over-reliance on single references and restrictive parametric assumptions about the data-generating process [11, 13, 15, 16]. These limitations are especially problematic for complex and dynamic tissue environments, such as cancer, where cells frequently exhibit transitional or hybrid states that are not represented in reference datasets. In addition, existing methods are often computationally inefficient and difficult to extend for spatial omics data beyond transcriptomics, such as spatial proteomics and epigenomics [17, 18].

To address these challenges, we present Φ-Space ST, a novel computational framework designed for platform-agnostic annotation of cell states in ST data. Φ-Space ST builds on Φ-Space, previously developed for single-cell multiomics analysis [19] (Fig 1A). Φ-Space ST leverages partial least squares (PLS) regression to annotate spatial cell-like objects with a continuous cell type scores. Unlike existing deconvolution methods, Φ-Space ST enables annotation using multiple scRNA-seq references, allowing the identification of both known and novel cell states. Its nonparametric and flexible framework avoids restrictive assumptions, offering greater adaptability. Additionally, Φ-Space ST is significantly faster than state-of-the-art methods like RCTD [11] and cell2location [13]. These features make Φ-Space ST highly versatile for a wide range of ST platforms and biological contexts.

**Fig 1:**
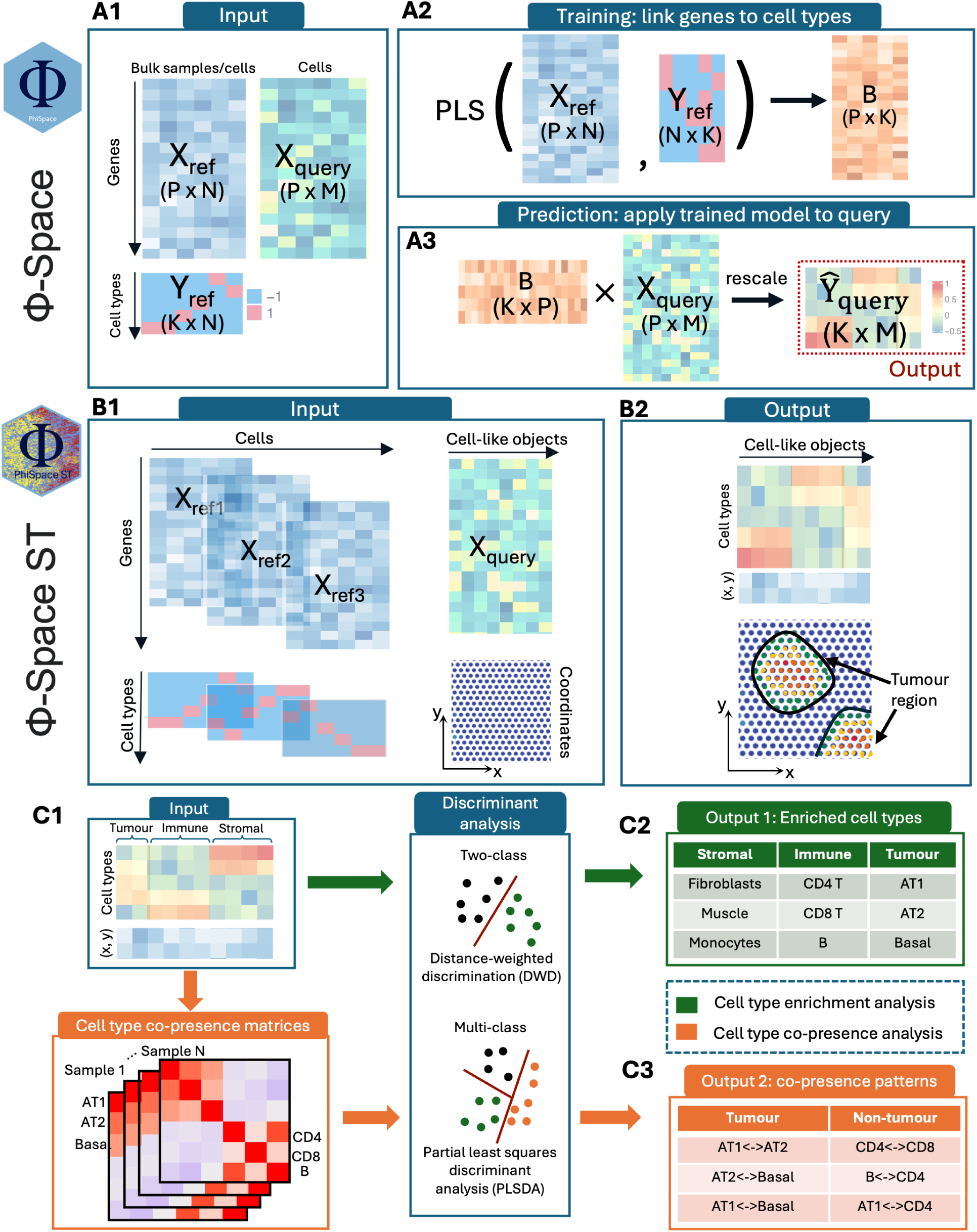
Schema of Φ-Space ST. **A:** Framework of the transcriptomic module of our previous Φ-Space method [19]. **A1:** Φ-Space enables continuous phenotyping of cells based on either bulk or single-cell RNA-seq references. **A2:** We first compute a partial least squares (PLS) regression coefficient matrix *B* to predict all *K* cell types on a continuous scale using *P* gene expression levels. **A2:** We then apply *B* to query gene expression matrix to obtain continuous cell type scores *Ŷ*_query_ for each of the query cells. **B:** Framework of Φ-Space ST, a new platform-agnostic and computationally efficient method designed to identify continuous cell states of spatial cell-like objects (supercellular spots, segmented cells or spatial bins) using spatial transcriptomics (ST) data. **B1:** We first train PLS regression models from one or multiple scRNA-seq references to continuously predict reference cell types. **B2:** We then apply the PLS models to the query ST data. Φ-Space ST accurately identifies not only the within-reference cell states, but also out-of-reference, e.g. transitional and malignant, cell states. **C:** Analyses based on Φ-Space ST continuous annotations of query cell-like objects, which we refer to as their *phenotype space embeddings*. **C1:** We identify spatial niches by clustering the phenotype space embeddings. **C2:** Cell type enrichment analysis, where we identify differentially enriched cell types in each spatial niche. **C3:** Cell type co-presence analysis, where we quantify the spatial co-presence of cell types and then use discriminant analysis to infer how specific co-presence patterns are associated with disease conditions.

In this study, we illustrate the application of Φ-Space ST to three cancer studies involving diverse ST platforms including 10x Genomics Visium [2], NanoString CosMx [3] and BGI’s Stereo-seq [4]. We demonstrate the ability of Φ-Space ST in identifying spatial niches, enriched cell types and cell type co-presence patterns that distinguish tumour from non-tumour tissue regions. By offering robust, scalable and biologically interpretable annotations, Φ-Space ST sets the stage for more comprehensive analyses of spatially resolved transcriptomics data.

## 2 Results

### 2.1 Φ-Space ST: a platform-agnostic method for identifying continuous cell states from ST data

#### Overview

Φ-Space ST is a platform-agnostic method for annotating different types of *cell-like objects* in ST data. We define a cell-like object as as the collection of transcripts observed in the proximity of a certain tissue location. Depending on the technical specifications of ST platforms, cell-like objects can be multi-cellular spots (e.g. 10x Visium [2]), segmented cells [8] or bins of aggregated transcripts (e.g. Stereo-seq [4]).

#### Multi-reference learning of cell states

Φ-Space ST transfers the rich knowledge about cell states from multiple reference scRNA-seq datasets to the query ST data, based on the assumption that all cell states in the query data can be characterised as mixtures of cell types defined in the reference, including out-of-reference cell states that may be present in the query. For example, some lung cancer cells may be jointly characterised by healthy lung epithelial *and* mesenchymal cell types [20].

#### Partial least squares as methodological backbone

The core of Φ-Space ST is partial least squares (PLS) regression [21, 22]. The first step of the Φ-Space ST workflow is to select genes that are useful for predicting cell types defined in the reference datasets. In the second step (Fig 1B1), a PLS regression model is trained from each reference scRNA-seq dataset to predict the reference cell type on a continuous scale. In the third step (Fig 1B2), the PLS models are applied to the query ST dataset. Each query cell-like object is characterised by its similarities with individual cell types defined in all references. We refer to this continuous annotation of cell-like objects as their *phenotype space embeddings*.

#### Quantifying cell type divergence in complex tissue microenvironment

Conventional cell type deconvolution methods characterise the cell state of a cell-like object as a *composition* of cell types in the reference, which requires the predicted cell type scores of each cell-like object to sum up to 1 [11, 13–16]. While this is a reasonable assumption in some scenarios, it may become stringent in disease samples such as the tumour microenvironment, where tumour and tumour-related cells typically exhibit divergent cell states [23]. The restriction of cell type scores to sum up to 1 can introduce compositional bias, which may hinder biological meaningful comparison of cell states across tissue regions and samples (e.g. the difference between transcriptionally active and inactive tissue regions may be masked) [13].

Considering the complex cell state landscape of disease-altered tissue microenvironment, Φ-Space ST removes the assumption that the query cell state can be characterised as a composition of reference cell types. As a result, we can measure the *cell state divergence* of a query cell-like object, defined as the 75% quantile of all predicted cell type scores. Higher cell state divergence implies that a cell-like object exhibits similarity to more reference cell types, flagging stronger transcriptional activity or transitional cell states.

#### Phenotype space embeddings for multi-sample biological discovery

The phenotype space embeddings of the cell-like objects from different tissue samples are in the same scale and hence provide a basis for multi-sample biological discoveries from ST data. We use phenotype space embeddings of cell-like objects to identify spatial niches, niche-specific enriched cell types and cell type co-presence patterns that separate disease from healthy samples (Fig 1C; details in Methods Section 4).

#### Summary of case studies

We apply Φ-Space ST to three cancer studies to showcase the biological insight provided by Φ-Space ST for identifying spatial phenotypic patterns in tumour microenvironments based on data from a diverse range of ST platforms. The case studies are summarised in Table 1 along with the annotation challenges faced for each of these platforms.

**Table 1:** Summary of case studies and associated analytical challenges. Challenge 1: Annotating tumour-associated cell states using healthy cell states; Challenge 2: Leveraging multiple scRNA-seq references; Challenge 3: Providing tnterpretable cell type deconvolution in tumour microenvironment; Challenge 4: Identifying spatial niches based on lineage tracing information. N: number of cells; NSCLC: non-small cell lung cancer; CITE-seq: cellular indexing of transcriptomes and epitopes by sequencing; AML: acute myeloid leukaemia.

| Platform | Reference | Query | Challenges |
| --- | --- | --- | --- |
| **10x Visium** [2]:<br>supercellular<br>sequencing-based<br>(whole-transcriptome) | Subsample ($N = 58,759$) of Human Lung Cell Atlas with 61 healthy cell types [27] | 18 Visium tissue samples from NSCLC patients and healthy donors [24].<br>*Cell-like objects*: multicellular spots (diameter 55 $\mu\text{m}$) | 1, 3 |
| **NanoString CosMx** [3]:<br>subcellular<br>imaging-based (using a gene panel of $\sim 980$ genes) | 4 scRNA-seq datasets from healthy and fibrotic lungs, each including cells from one of the following lineages: immune, epithelial, endothelial and mesenchymal [28] | 8 CosMx tissue samples from healthy and NSCLC donors [25].<br>*Cell-like objects*: segmented cells | 1, 2 |
| **BGI Stereo-seq** [4]:<br>subcellular<br>sequencing-based<br>(whole-transcriptome) | 4 scRNA-seq datasets: Spleen, mouse spleen [29]; BM, mouse bone marrow [30]; Neutro, neutrophils from healthy and cancerous mice [31]; CITE, CITE-seq (only used the RNA part) of mouse spleen immune cells [32] | 1 Stereo-seq mouse spleen sample with from an AML mouse with cellular lineage tracing barcodes.<br>*Cell-like objects*: spatial bins ('bin50', i.e. sidelength 25 $\mu\text{m}$) | 1–4 |

We explain in detail the different characteristics of ST data analysed in these case studies:

- *10x Visium* [2] is a sequencing-based platform, which provides whole-transcriptome sequencing in spatial spots with diameter 55 µm. Each spot may contain multiple cells and hence can be viewed as a mini-bulk. The Visium dataset we analysed consists of 18 lung sections selected from 9 individuals, including 1 healthy donor and 8 non-small cell lung cancer (NSCLC) patients (that is, two sections from each patient) [24].
- *Nanostring CosMx* [3] is an imaging-based platform, which is capable of detecting transcripts with subcellular precision. The CosMx dataset we analysed consists of 8 tumorous lung tissue sections from 5 NSCLC patients [25]. The expression level of 980 genes were measured on each tissue sample. We used the morphology-based cell segmentation results provided by He et al. [25] and analysed transcriptional profiles of segmented cells.
- *BGI Stereo-seq* [4] is a sequencing-based platform, which provides whole-transcriptome sequencing at subcellular resolution. The Stereo-seq dataset we analysed consists of one mouse spleen section containing acute myeloid leukaemia (AML) cells [26]. Singlecell profiling and lineage tracing (SPLINTR) was used to barcode the AML cells, so that descendants from the same ancestor AML cell (i.e. an AML clone) shared the same barcode. Due to a technical issue known as ‘spot bleeding’ [10], no reliable cell segmentation could be conducted to infer cell boundaries. Hence, following Holze et al. [26], we binned the transcripts into ‘bin50’, i.e. spatial bins of sidelengths ∼25 µm, and analysed these spatial bins.

### 2.2 Φ-Space ST provides highly interpretable cell type deconvolution for understanding cell type co-presence

For supercellular spatial platforms such as 10x Visium, single-cell resolution is technically unattainable (see Section 2.1). As a result, to infer spatial distribution of cell types, deconvolution algorithms must be used to estimate the abundance of different cell types within each multicellular spot. In this case study, we first demonstrate that Φ-Space ST provides more interpretable cell type deconvolution compared to some state-of-the-art deconvolution methods and is very fast to compute. We then use the Φ-Space ST deconvolution results to infer cell type co-presence patterns. This analysis leads to an interpretable separation of samples from different subtypes of non-small cell lung cancer (NSCLC), revealing potential connections between cell type co-presence and NSCLC prognosis.

#### Φ-Space ST provides more interpretable cell type deconvolution

In this benchmark we applied Φ-Space ST, RCTD [11], cell2location [13] and TACCO [16] to the Visium NSCLC dataset [24]. These four methods produced discrepant estimates of the spatial distribution of cell types. For example, for a tumorous lung section P11 T3 (Fig 2A), the B cell abundance in spatial spots inferred by these methods might lead to very different biological interpretations on how B cells infiltrated the tumour region (Fig 2B). Given the important role of B cells in tumour microenvironment [33, 34], these diverging estimates may result in very different cancer prognosis and treatment.

**Fig 2:**
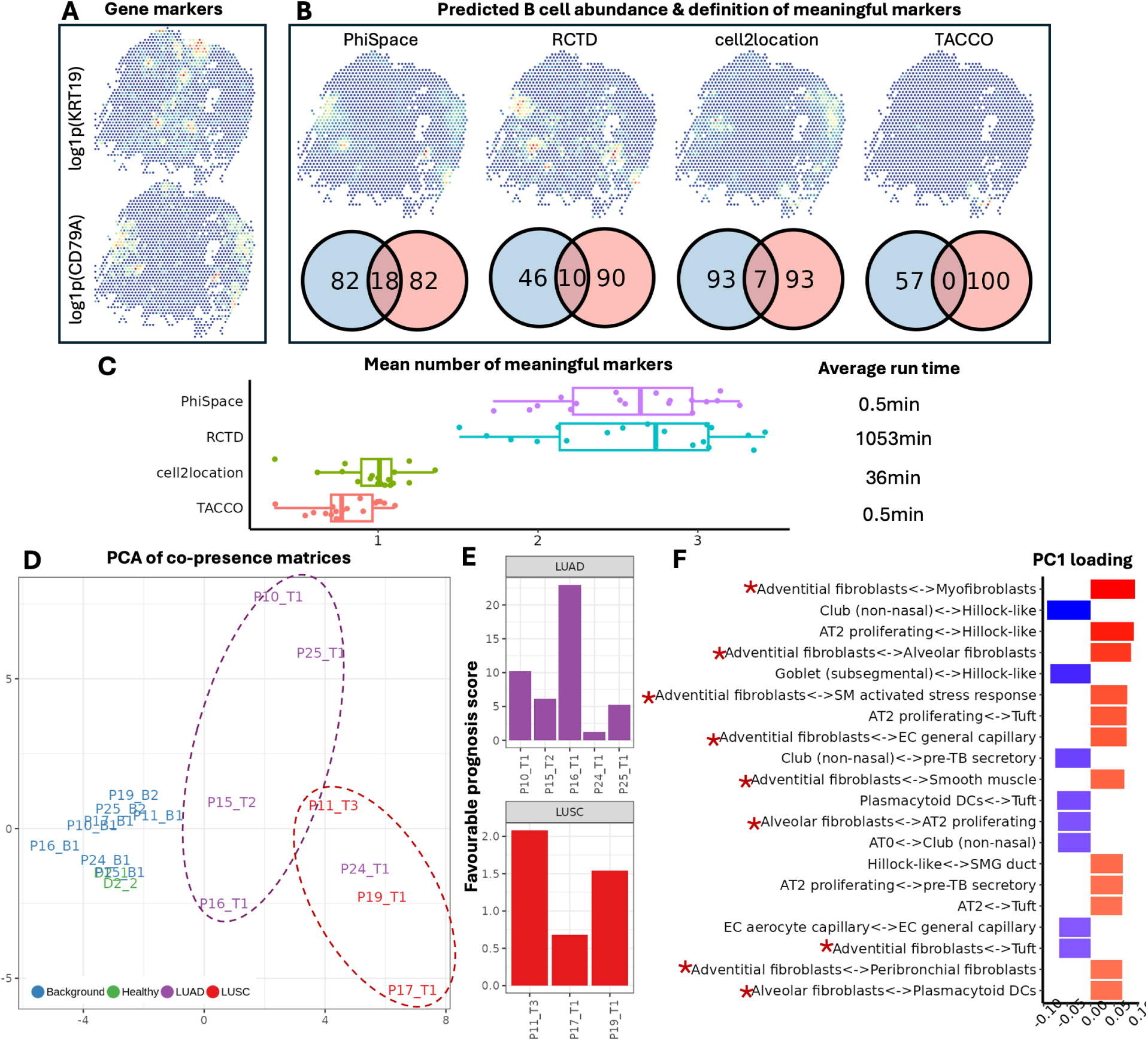
Benchmarking of cell type deconvolution methods using Visium lung cancer samples. **A:** A 10x Visium non-small cell lung cancer (NSCLC) tissue P11 T3 coloured according to expression levels of NSCLC marker KRT19 and B cell marker CD79A, where each dot represents a 55 µm multicellular spot. **B:** Predicted B cell abundance in P11 T3 according to Φ-Space ST and three state-of-the-art cell type deconvolution methods. These four methods resulted in very different estimates of B cell spatial distribution. To evaluate the quality of B cell abundance predictions, we calculated the number of meaningful markers for B cell prediction by each method: in each Venn diagram, the blue circle represents differentially expressed genes in the spatial region predicted to have high B cell abundance; the red circle represents B cell marker genes learned from the reference. **C:** Boxplot comparing mean number of meaningful markers per tissue for all 18 tissues. Average run time of each method is also shown. **D:** We computed cell type co-presence matrices for 18 lung tissues and computed their first two principal components (PCs) with proportions of variance explained by these PCs. The PC plot shows a clear separation between non-tumorous (healthy and background) and tumorous (LUAD and LUSC) samples. Furthermore, LUSC, which tend to have worse prognosis than LUAD, tended to be separated from LUAD.(LUAD, lung adenocarcinoma; LUSC, lung squamous cell carcinoma; background, non-tumorous tissues from cancerous lungs.) **E:** To investigate if the heterogeneity of tumorous samples observed in **D** reflected differential prognoses, we computed the favourable prognosis scores for both LUAD and LUSC samples. P24 T1 had the least favourable predicted prognosis among the LUAD samples, and was located closer to LUSC samples in **D**; P11 T3 had the most favourable predicted prognosis among the LUSC samples, and was located closer to LUAD samples. In general, tumorous samples tended to form two clusters (circled in **D**) according to prognoses. **F:** Since different disease conditions were mainly separated along PC1 in **D**, we plotted the top 20 cell type co-presence patterns contributing to PC1: 9/20 involved fibroblasts (marked by ⋆).

Evaluating the accuracy of cell type deconvolution is challenging due to the lack of available ground truth. Thus we defined *meaningful markers* of a target cell type in a given tissue sample as genes in the intersection of two gene sets: the differentially expressed marker genes learned from the reference data, and the spatially differentially expressed genes in the query tissue area with high predicted abundance of the target cell type (see Supplementary Methods Section S2). The number of meaningful markers measures how cell type deconvolution matched prior knowledge about cell types via marker gene identification. When predicting B cell abundance in the tissue section P11 T3, Φ-Space ST had the highest number of meaningful markers (Fig 2B).

Then, to evaluate the performance of the deconvolution methods on all cell types for tissue P11 T3, we calculated the mean number of meaningful markers averaged across all 61 cell types defined in the reference (Supplementary Fig S1A). Φ-Space ST had the highest mean number of meaningful markers. Finally, across all 18 tissue samples (Fig 2C), Φ-Space ST and RCTD had significantly higher mean number of meaningful markers compared to cell2location and TACCO. In addition, Φ-Space ST only took less than 1 minute to compute each tissue sample, much faster than RCTD, which took hours to compute each tissue sample.

#### Φ-Space ST reveals lung cancer subtype-specific cell type co-presence

With high throughput and multicellular spots, Visium samples have been viewed as particular suitable for investigating cell type co-presence [35, 36]. A key to biologically meaningful co-presence analysis is reliable cell type deconvolution, a task that Φ-Space ST excels at (Fig 2A–C). Therefore, we computed the cell type co-presence matrices based on cell type abundance inferred by Φ-Space ST for the Visium samples. We computed one cell type co-presence matrix for each of the 18 Visium samples, then extracted the principle components (PCs) of the co-presence matrices (details in Methods Section 4).

We observed a clear separation between LUAD, LUSC and non-tumorous samples (Fig 2D). Moreover, the LUAD and LUSC samples appeared to be more heterogeneous compared to the non-tumorous samples, with LUAD samples and LUSC samples separately grouped to-gether. Since LUSC tends to have worse prognosis than LUAD [37], we investigated whether the observed separation might be due to the differential prognoses of cancer patients. As we lacked additional patient-level information from the original publication [24], we opted for a computational approach to predict prognosis based on markers identified in the Visium samples [37] (see Supplementary Methods Section S2). Our analysis identified LUAD (LUSC) samples with the least (most) favourable prognosis (Fig 2E). This was confirmed by the location of sample P24 T1 (least favourable prognosis) close to the LUSC samples and P11 T3 (most favourable prognosis) close to the LUAD samples (Fig 2D). Therefore we were able to attribute the heterogeneity of tumorous samples observed on Fig 2D to their differential prognoses.

Since the disease conditions were mainly separated along the first principal component (PC1) on Fig 2D, we investigated how co-presence of specific pairs of cell types contributed to this separation by examining the top 20 PC1 cell type co-presence loadings (Fig 2F). Notably, adventitial and alveolar fibroblasts were featured in 9 of the top 20 loadings, highlighting the diverse roles of fibroblasts in lung tumour microenvironment [38, 39]. To focus on solely on immune cell types, we excluded non-immune cell types in the co-presence matrices and reran the analysis. We observed that the separation between LUAD, LUSC and non-tumorous tissues was largely retained (Supplementary Fig S1B), with plasmatoid dendritic cell (pDC) playing a prominant role in explaining this separation (Supplementary Fig S1C) [40]. In particular, the co-presence of pDC and different types of fibroblasts all had positive loadings, suggesting that the pDC–fibroblast co-presence might be related to lung cancer prognosis [41].

Deconvolution results from other methods can also be used to conduct the cell type co-presence analysis featured in Fig 2D–F. However results based on alternative methods were less interpretable compared to Φ-Space ST. For example, although the first two PCs of co-presence matrices based on RCTD also showed a good separation of disease conditions, the PCs there lacked a correspondence to cancer prognosis as in the Φ-Space ST results above (Supplementary Fig S2).

#### Summary

Through the Visium case study we showed that when single-cell resolution was technically unattainable, Φ-Space ST provided biologically interpretable cell type deconvolution results. Compared to state-of-the-art methods specifically designed for cell type deconvolution, Φ-Space ST deconvolution either led to increased interpretability or faster computation. A multi-sample cell type co-presence analysis enabled to identify differential cell type co-presence patterns in different cancer subtypes, suggesting some potentially association between cell type co-presence and cancer prognosis.

### 2.3 Φ-Space ST characterises heterogeneity of lung tumour microenvironment using healthy single-cell reference atlas

Φ-Space ST can uncover complex cell states in tumour microenvironment by combining multiple scRNA-seq references, as we shall demonstrate below using the CosMx dataset from NSCLC patients (detailed in Section 2.1). Since cell segmentation was available for these tissue sections [25], we treated segmented cells as cell-like objects. We obtained biologically meaningful identification of altered cell states by leveraging multiple scRNA-seq references from healthy and fibrotic lungs, highlighting the ability of Φ-Space ST in identifying divergent cell states.

#### Φ-Space ST uncovers complex and heterogeneous cell state landscapes of tumour microenvironment

We first focused on a particular lung section Lung5 Rep1 from a lung adenocarcinoma (LUAD) patient to demonstrate how Φ-Space ST can provide insights into the cell state heterogeneity of the tumour microenvironment. Nine spatial domains were identified in this tissue section by He et al. [25] (Fig 3A). According to our Φ-Space ST annotation, the tumour interior, tumour-stroma boundary and lymphoid structure domains had higher cell type divergence (Supplementary Fig S3; see also Methods Section 4). The lymphoid structures were highly divergent, potentially due to the dense population of various immune cell types, such as T, B and dendritic cells [42] as identified by Φ-Space ST (Fig 3B). The tumour domains were highly divergent, consistent with lung tumours tending to exhibit molecular features of multiple cell types [37]. We found that a combination of mesenchymal, epithelial and immune cell types were enriched in the tumour domains, including tumour interior and tumour-stroma boundary (Fig 3B). In particular, the secretory and transitional AT2 cell types reflected the glandular cell origin of LUAD [43]; the observed basal and KRT5-/KRT17+ epithelial cell types in LUAD was consistent with previous study [44]; and the presence of mesothelial and activated myofibroblast cell types flagged the potential existence of epithelial-to-mesenchymal transition [45] and cancer-associated fibroblasts [46].

**Fig 3:**
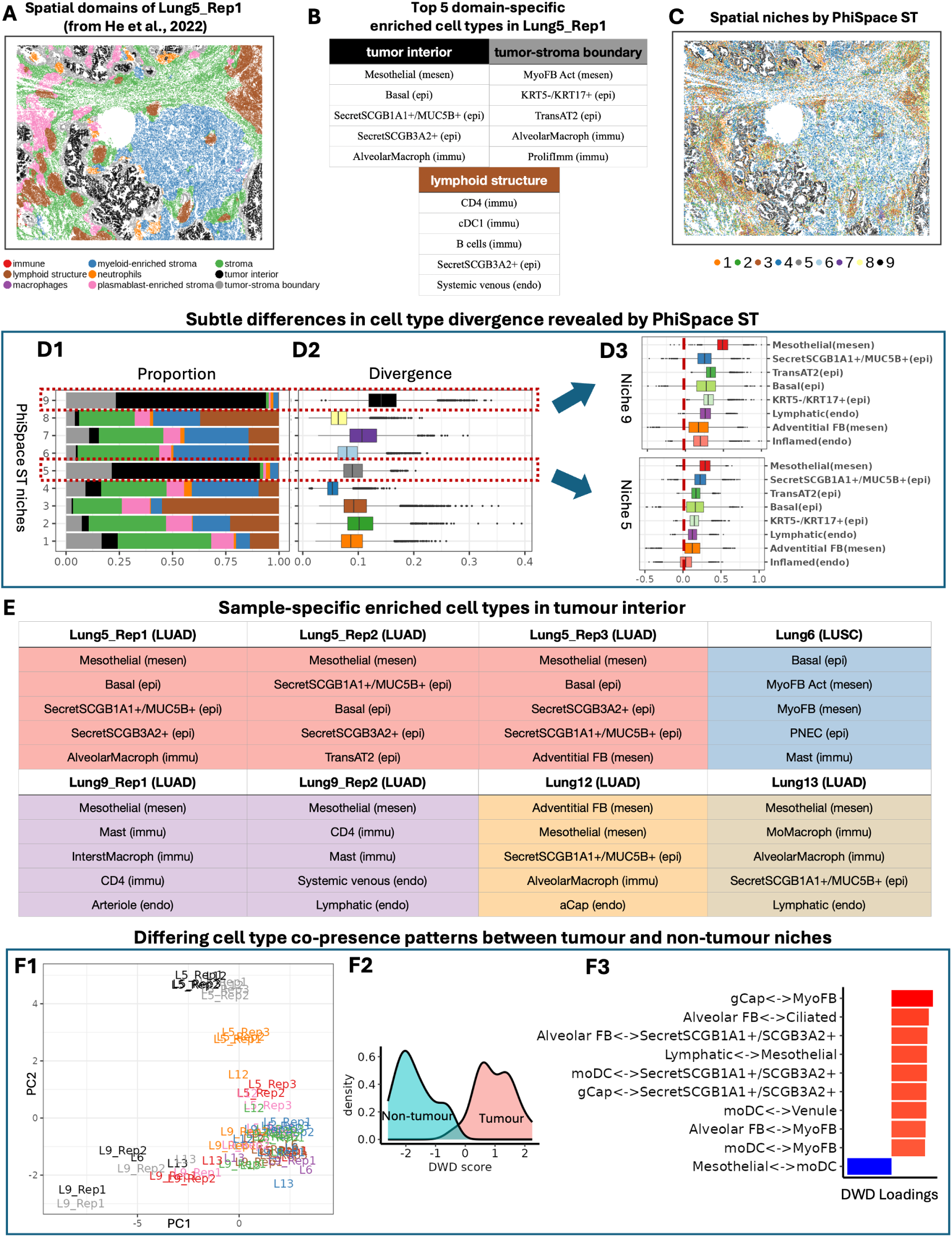
Exploring cell state heterogeneity in CosMx lung samples. **A:** CosMx lung sample (Lung5 Rep1) from a patient with lung adenocarcinoma (LUAD), with spatial domains identified by He et al. [25]. Each dot represents the location of the nucleus of a cell. **B:** Top 5 most enriched cell types of some spatial domains in the sample Lung5 Rep1 identified by Φ-Space ST. (SecretSCGB1A1+/MUC5B+: secretory cell with unregulated SCGB1A1 and MUC5B; SecretSCGB3A2+: secretory cell with upregulated SCGB3A2; AlveolarMacroph: alverolar macrophage; MyoFB Act: activated myofibroblast; TransAT2: transitional type-2 alveolar cell; ProlifImm: proliferating immune cell; CD4: CD4 T cell; cDC1: type-1 conventional dendritic cell.) **C:** Φ-Space ST niches, identified by clustering phenotype space embeddings of segmented cells. **D:** Cell type divergence of Φ-Space ST niches. **D1:** Cell proportions according to spatial domains (**A**) for each Φ-Space ST niche. Niches 5 and 9 contained large proportions of cells from tumour domains (tumour interior and tumour-stroma boundary). **D2:** Cell type divergence of Φ-Space ST niches. The two tumour niches 5 and 9 differ greatly in their divergence. **D3:** Φ-Space ST scores of the most enriched cell types in niches 5 and 9. While these two niches had similar enriched cell types, the scores in niche 9 were generally higher than in niche 5, leading to the difference in cell type divergence. **E:** Top 5 most enriched cell types in tumour interior domains of different lung samples. (LUAD: lung adenocarcinoma; LUSC: lung squamous cell carcinoma.) **F:** Cell type co-presence analysis. **F1:** First two principal components (PCs) of all the domain-specific co-presence matrices. Tumour domains (tumour interior and tumour-stroma boundary) and non-tumour domains were separated, reflecting distinct cell type co-presence patterns. **F2:** Discriminant analysis (DWD) of all domain-specific co-presence matrices shows clear separation between tumour and non-tumour domains. **F3:** DWD loadings highlighting most discriminant co-presence patterns, where two cell types having positive (negative) loading indicates if their co-presence contributes to the prediction of tumour (non-tumour) domains. (gCap: general capillary cell; FB: fibroblast; Secret: secretory cell; moDC: monocyte-derived dendritic cell.)

#### Φ-Space ST identifies heterogeneous tumour niches with differential cell type divergence

To compare Φ-Space ST spatial niches with the spatial domains identified by He et al. [25] (Fig 3A), we identified 9 spatial niches of Lung5 Rep1 based on the Φ-Space ST annotation (Fig 3C; see Methods Section 4). Overall, our Φ-Space ST niches corresponded well to the domains in Fig 3A. A notable difference was that we identified two tumour niches, niches 5 and 9, which both contained a large proportion of cells from the tumour interior and tumour-stroma boundary domains (Fig 3D1), but had significantly different cell state divergence (Fig 3D2). In particular, these two niches were enriched by the same cell types but niche 9 showed a much higher divergence compared to niche 5 (Fig 3D2&D3). This highlighted intra-tumour heterogeneity that was not captured in He et al. [25].

#### Φ-Space ST identifies differentially enriched cell types in different cancerous lung samples

As we have previously demonstrated [19], the phenotype space embeddings of query datasets derived using PLS regression models are robust against batch effects. This property of PLS also makes Φ-Space ST annotations from different query ST samples directly comparable. For example, the enriched cell types in the tumour interior domain were reproducible across biological replicates, i.e. tissue sections from the same lungs (see replicates from Lung5 and Lung9 in Fig 3E). In contrast, the enriched cell types in the tumour interior domain from different patients exhibited significant heterogeneity (compare Lung5, Lung6, Lung9, Lung12 and Lung13 in Fig 3E). Specifically, mesothelial cell type was enriched in all lung adenocarcinoma (LUAD) tumours (Lung5, 9, 12 and 13) but not in lung squamous cell carcinoma (LUSC) sample (Lung6). Two myofibroblast cell types were enriched in the LUSC sample (Lung6), consistent with a recent finding that LUSC tumours tended to have higher level of myofibroblasts compared to LUAD [47].

#### Φ-Space ST reveals cancer-specific cell type co-presence patterns

To investigate how cell type co-presence patterns distinguish spatial niches across samples, we calculated the cell type co-presence matrices for spatial domains in all 8 CosMx NSCLC samples, visualised using PCA (detailed in Methods Section 4). We observed a separation of tumour domains from non-tumour domains (Fig 3F1). Furthermore, a discriminant analysis using DWD (Fig 1C) led to an improved separation between tumour and non-tumour domains (Fig 3F2). The DWD loadings highlighted pairs of cell types whose co-presence in the same segmented cell contribute to the prediction of tumour or non-tumour domains (Fig 3F3). The tumour domains were characterised by complex interactions between endothelial (gCap, lymphatic, venule), epithelial (ciliated, secretory) and mesenchymal (myoFB, alveolar FB, mesothelial) cell types, indicating possible epithelial-mesenchymal, mesenchymal-epithelial and endothelial-mesenchymal transitions as typically seen in malignant tumours [20, 48]. In addition, the role of monocyte-derived dentritic cells (moDC) was also highlighted. The co-presence of moDC, secretory, venule and myofibroblast cell types in tumour domains was consistent with a previous observation that myeloid DCs (including moDC) were commonly observed in NSCLC specimens [49]; however, the co-presence of moDC and mesothelial tended to be observed in non-tumour domains, which may reflect the shared role of mesothelial cells and moDCs in antigen presentation [50].

#### Summary

Through the CosMx case study, we demonstrated that Φ-Space ST could robustly identify cancerous cell states based on four scRNA-seq multi-reference datasets from healthy and fibrotic lungs. We identified domain-specific enriched cell types that were biologically interpretable and comparable across different samples and tissue sections. Our cell type co-presence analysis identified patterns that separated tumorous from non-tumorous domains, highlighting how complex interactions between epithelial, endothelial, mesenchymal and immune cells have jointly shaped the lung tumour microenvironment.

### 2.4 Φ-Space ST achieves segmentation-free annotation of subcellular ST data

Clonally resolved ST data links spatial information with clonal identity. This technological advance promises to uncover transcriptional mechanisms that contribute to intratumour heterogeneity of cancer clones [26, 51]. However, ST data combined with lineage tracing remain rare and under-explored, with no established workflow. To address this challenge, we developed a Φ-Space ST-based workflow to provide unique spatial insights by incorporating information from cancer lineage tracing (Fig 4A). We investigated the heterogeneity of acute myeloid leukaemia (AML) clones based on a Stereo-seq mouse spleen dataset [26] (details in Section 2.1). Liquid tumours such as AML tumours have less clearer spatial structures compared to solid tumours [52], which adds to the challenges in analysing this dataset.

**Fig 4:**
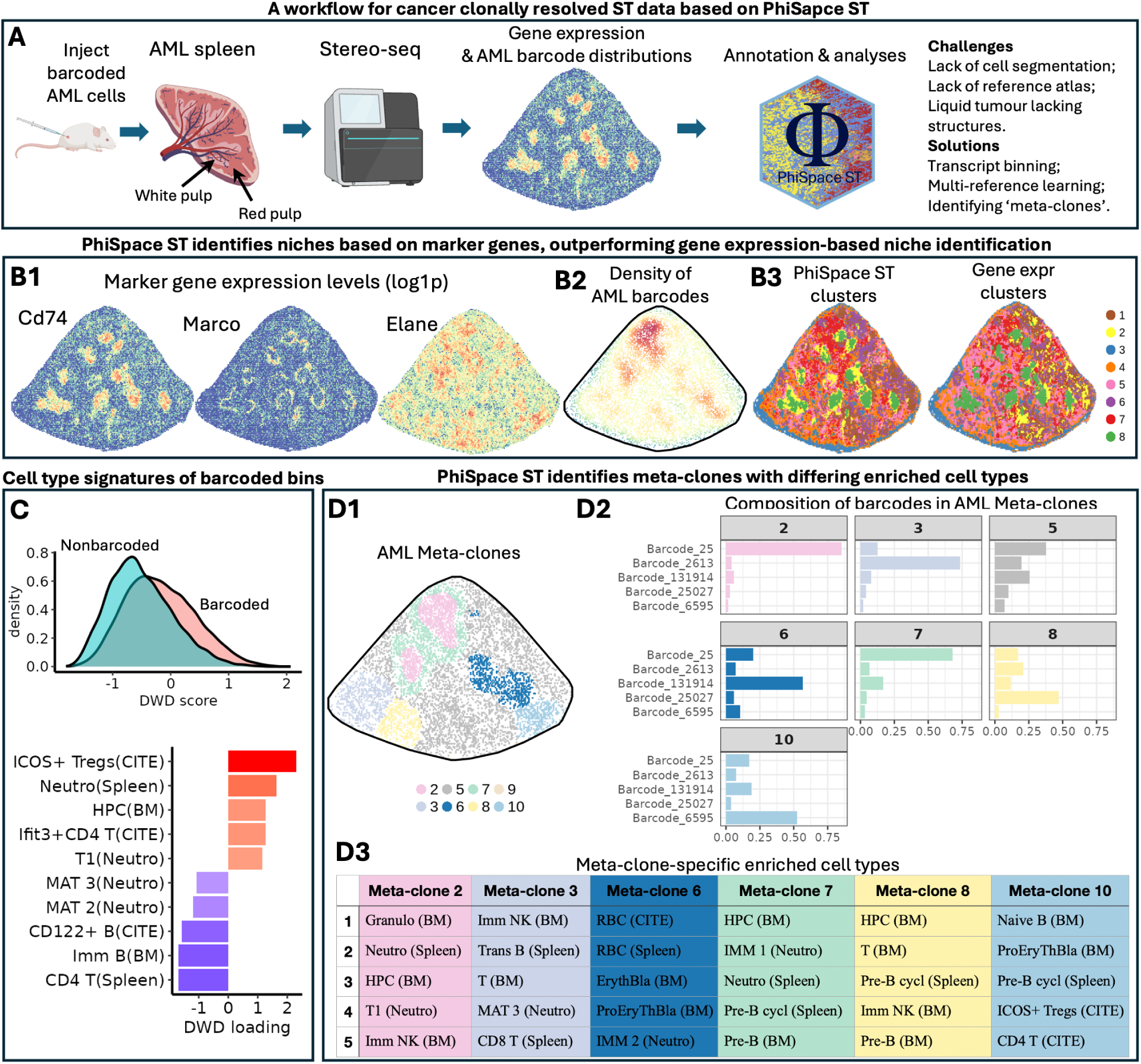
Uncovering cancer clone-specific cell states in Stereo-seq AML mouse spleen data. **A:** Overview of the workflow for analysing clonally-resolved ST data based on Φ-Space ST. Stereo-seq AML mouse spleen data with lineage tracing information were obtained from an AML mouse with barcoded cancer cells. (Created in https://BioRender.com.) Using gene expression and barcode information as input, Φ-Space ST identifies AML ‘meta-clones’ with differential enriched cell types based on multiple reference datasets. **B:** Φ-Space ST identifies more nuanced spatial niches compared to conventional gene expression-based niche identification. **B1:** Spatial distribution of morphological gene markers Cd74 (white pulp), Marco (marginal zone) and Elane (neutrophil). Each dot represents a spatial bin of sidelengths ∼25 µm where at least one transcript was detected. **B2:** Density of all AML barcodes in the mouse spleen sample. **B3:** Eight clusters of spatial bins were identified based on either Φ-Space ST phenotype space embeddings or gene expressions of the bins. The Φ-Space ST clusters better delineated marginal zone and the high barcode density regions shown in **A**2 compared to the gene expression clusters. **C:** A DWD discriminant analysis revealed differences between bins that contained AML barcodes (‘barcoded’) and bins that did not contain any barcodes (‘non-barcoded’). Barcoded bins tended to have larger DWD scores. Top 10 DWD loadings (ranked by absolute value) show most important cell types for distinguishing barcoded and non-barcoded bins. (CITE, Spleen, BM, Neutro: four reference datasets; Treg: regulatory T cell; Neutro: neutrophil; HPC: hematopoietic progenitor cell; T1: tumour-associated neutrophil; Mat 2, 3: mature neutrophil; Imm B: immature B cell.) **D:** AML meta-clones and their enriched cell types. **D1:** Eight meta-clones were identified based on their barcode compositions. Bins with similar barcode (clonal) compositions were grouped into the same meta-clone. **D2:** Proportion of 5 most abundant barcodes in each meta-clone (with the exception of meta-clone 9 that is too rare). The meta-clones had a diverse composition of individual AML clones, where all meta-clones, except 5, had a dominating AML clone. **D3:** Enriched cell types within individual meta-clones. Compared to **F**, neighbouring meta-clones (i.e. 2&7, 6&10 and 3&8) tended to have similar enriched cell types, whereas non-neighbouring meta-clones tended to have dissimilar enriched cell types (Granulo: granulocyte, including neutrophil; IMM NK: immature natural killer; Trans B: transitional B cell; RBC: red blood cell; (Pro)ErythBla: (pro)erythroblast; Imm 1, 2: immature neutrophil; Pre-B (cycl): (cycling) pre-B cell).

This Stereo-seq dataset lacks reliable cell segmentation, so we binned the whole tissue slide into bins of sidelengths ∼25 µm (as in Holze et al. [26]). Therefore cell-like objects in this case study were spatial bins. Since a comprehensive mouse spleen cell atlas is not available, we used Φ-Space ST to leverage multiple scRNA-seq reference datasets without the need for prior integration (details in Methods Section 4). Since we wished to make use of a scRNA-seq dataset generated from the same mouse spleen as the Stereo-seq sample [51], we used a ‘bridging’ strategy for continuous annotation transfer with Φ-Space ST. Our approach is very different from other deconvolution methods which cannot transfer continuous annotations. We first annotated the intermediate scRNA-seq spleen sample, then used the scRNA-seq spleen sample (with continuous Φ-Space ST annotations) to annotate the Stereo-seq sample.

#### Φ-Space ST multi-reference annotation provides a comprehensive characterisation of AML spleen

The mouse spleen has a distinctive morphology, where T and B lymphocytes in the white pulp coordinate adaptive immune responses. Surrounding the white pulp is the marginal zone, a region enriched with macrophages, dendritic cells and marginal zone B cells that capture antigens and facilitate their delivery to lymphocytes in the white pulp [53] (Figure 4B1). Specific to this spleen sample was the prevalence of Elane expression (a neutrophil marker). As lineage tracing information was available we were able to locate the AML cells (Fig 4B2).

The four reference datasets we used contained immune cell types from different mouse tissue types, including mouse spleen, bone marrow (origin of the myeloid lineages of immune cells) and tumour (details in Section 2.1). These references jointly provide a comprehensive characterisation of the immune landscape in AML mouse spleen. We first clustered the spatial bins based on their multi-reference phenotype space embeddings, which resulted in more subtle spatial regions compared to the clusters based on gene expression (Figs 4B3; see Supplementary Methods Section S2). Thus our Φ-Space ST approach successfully captured biologically meaningful spatial variations across morphological structures in the spleen.

We further evaluated whether the Φ-Space ST cell type scores could reveal the differences between bins that contained AML barcodes and non-barcoded bins. Our DWD analysis (detailed in Methods Section 4) showed that barcoded bins tended to have stronger ICOS+ regulatory T cell (ICOS+ Treg), red blood cell (RBC), neutrophil (Neutro, T1), hematopoietic progenitor cell (HPC) and CD4 T cell identities, compared to non-barcoded bins (Fig 4C). The strong ICOS+ Treg identity of barcoded bins likely reflects the ability of AML cells to promote the expansion of Tregs by expressing ICOS ligand [54]. The neutrophil identity of the barcoded region may be explained by the high expression of the neutrophil maker Elane in that region (Fig 4B1). Notably, the barcoded and non-barcoded bins were enriched by different types of neutrophils, with barcoded bins enriched by tumour-associated T1 neutrophils and non-barcoded bins enriched by mature neutrophils (MAT2, MAT3).

#### AML clones form phenotypically distinct meta-clones

In this clonally resolved Stereo-seq dataset, we observed that individual AML clones in mouse spleen tended to be concentrated in specific spatial domains, which motivated us to define ‘meta-clones’, i.e. spatial niches of bins with similar composition of AML clones (Supplementary Fig S4). We aggregated the spatial bins into meta-clones according to their clonal compositions and identified 8 meta-clones with significantly different compositions of individual AML clones (Fig 4D1&D2; details in Supplementary Methods Section S2).

The AML meta-clones exhibited differential cell states, characterised by their most enriched cell types compared to non-barcoded bins (Fig 4D3; details in supplementary Methods Section S2). We observed that meta-clone 7 formed the ‘periphery’ of meta-clone 2. Both meta-clones had strong neutrophil identities. The neighbouring meta-clones 6 and 10 both had strong erythroid identities while the neighbouring meta-clones 3 and 8 both had strong lymphoid identities (Figs 4D2&D3). In addition, non-neighbouring meta-clones tended to have enriched cell types from different lineages. These observations provided unique insights into the spatial organisation of AML tumours and their heterogeneity in terms of cell states, which remain poorly understood.

#### Summary

We showed how Φ-Space ST can facilitate biological discoveries in Stereo-seq data in a case where cell segmentation was not attainable, and where a suitable reference atlas was lacking. This case study focusing on liquid tumours also highlighted the challenge of tumour niches that are difficult to define. We used four scRNA-seq references to derive the Φ-Space ST phenotype space embeddings of Stereo-seq bins, which retained the complex morphological information of a cancerous spleen, and highlighted the cell states of AML bins. Our novel ways of defining meta-clones and identifying meta-clone-specific enriched cell type provide a generic approach for investigating cancer clonal heterogeneity using clonally resolved ST data.

## 3 Discussion

Φ-Space ST addresses key challenges in annotating cell states in complex tissues and disease environments. By leveraging the power of partial least squares (PLS) regression to annotate cell-like objects on a continuous scale using multiple references, Φ-Space ST provides a unified, platform-agnostic and computationally efficient framework for analysing ST data at multiple resolutions. The continuous cell state annotation enables the characterisation of both known and novel cell types, overcoming limitations of traditional methods that rely on single reference and are computationally costly.

Through three diverse case studies, we demonstrated the versatility and effectiveness of Φ-Space ST in unravelling the cellular and spatial complexity of tumour microenvironments. In the Visium case study, Φ-Space ST provided fast and interpretable deconvolution results, revealing cell type co-presence patterns associated with different cancer subtypes and prognostic outcomes. In the CosMx case study, Φ-Space ST’s continuous annotation strategy uncovered intra- and inter-tumour heterogeneity, highlighting distinct spatial niches and niche-specific enriched cell types. Finally, in the Stereo-seq case study, Φ-Space ST successfully characterised AML clone heterogeneity without requiring cell segmentation, showcasing its robustness in cases where traditional approaches fail.

Φ-Space ST’s innovative continuous annotation approach goes beyond static classification to offer rich insights into transitional and hybrid cell states, particularly in disease contexts such as cancer. Φ-Space is a highly flexible and computationally efficient tool for largescale and high-resolution spatial studies that provides a scalable solution to the increasing demands of modern spatial omics data.

Future work will focus on extending Φ-Space ST to other pathological conditions and integrating additional omics assays, such as spatial proteomics and epigenomics, to further expand its applicability. Validating its performance in larger and even more diverse datasets will also enhance its generalisability and utility across a wider range of biological contexts.

## 4 Methods

We denote the *R* reference datasets by (*X*_ref*,r*_*, Y*_ref*,r*_), *r* = 1*, …, R*, where each *X*_ref*,r*_ is a gene by cell scRNA-seq gene expression matrix of normalised values (Fig 1A). The phenotype matrix *Y*_ref*,r*_ is a dummy matrix that represents the *K_r_*cell type labels of cells in *X*_ref*,r*_. That is, each row of *Y*_ref*,r*_ represents a cell type and a cell is assigned the value 1 if it belongs to that cell type or −1 otherwise (Fig 1A).

We denote the query ST dataset by (*X*_query_*, S*_query_), where *X*_query_ is the query gene expression matrix with rows representing genes and columns representing spatial cell-like objects (segmented cells, multicellular spots or transcript bins), and where *S*_query_ contains the coordinates of the cell-like objects (*x, y*).

### 4.1 Φ-Space ST multi-reference continuous phenotyping

The main steps for deriving the Φ-Space ST cell type scores based on multiple references are as follows:

1. **Gene selection**. For each reference dataset (*X*_ref*,r*_*, Y*_ref*,r*_), we use the variable selection method based on partial least squares (PLS) regression, as described in Mao et al. [19], to select the most useful genes G*_r_* for predicting the cell type *Y*_ref*,r*_. That is, each G*_r_* contains the genes with the largest absolute values of PLS regression coefficients for predicting *Y*_ref*,r*_.
2. **PLS model training** (Fig 1B1). For the *r*th reference, we train a PLS regression model [21, 22] using only genes shared by G*_r_*, the selected genes for reference *r*, and *G*, all genes sequenced in the query. We estimate a low-rank regression coefficient matrix *B_r_*of size (|G*_r_*| × *K_r_*) to fit the regression model *Y*_ref*,r*_ = *X̃*_ref_*_,r_B_r_*, where *K_r_* denotes the number of cell types in the *r*th reference and *X̃*_ref_*_,r_* denotes the *r*th gene expression matrix using only selected genes. The rank of *B_r_* is equal to the number of PLS components, by default set to *K_r_*.
3. **Prediction** (Fig 1B2). We compute the continuous predicted cell type scores *Ŷ*_query_*_,r_* as a rescaled version of *X̃*_query_*B_r_*, where *X̃*_query_ denotes the query gene expression matrix using only selected genes; see Mao et al. [19] for details of the rescaling. The final predicted cell type scores *Ŷ*_query_ is the concatenation of all *Ŷ*_query_*_,r_* and has size 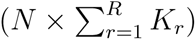, where *N* is the number of cell-like objects in the query. That is, the number of columns of *Ŷ*_query_ is equal to the sum of cell types defined in all *R* references.

We call the predicted cell type score matrix *Ŷ*_query_ the *phenotype space embeddings* of the query cell-like objects. Compared to hard cell type classification of cell type which assigns a unique cell type label to each query cell-like object, the phenotype space embedding contains much richer information regarding the spatial distribution of cell states and can be used for various downstream analyses described below.

### 4.2 Multi-sample spatial biological discovery

Φ-Space ST is a novel method that uses the phenotype space embeddings *Ŷ*_query_ to facilitate multi-sample biological discoveries in ST data.

#### Spatial niche identification (Fig 1C1)

Φ-Space ST provides a simple approach for spatial niche identification, a major analytical task in ST data analysis to partition the whole spatial domain into cell communities [9, 55]. We conduct k-means clustering based on the PCs of the Φ-Space ST cell type scores, where the number of PCs is selected via scree plot. Using PCs rather than the original cell type scores allows to denoise some cell type scores that may not be spatially variable enough to contribute towards niche identification.

#### Niche-specific enriched cell types (Fig 1C2)

Once spatial niches have been identified, we further the analysis to identify niche-specific enriched cell types. We perform discriminant analysis using one of the following two methods:

- *Distance-weighted discrimination (DWD) for two niches* [56]. DWD is a powerful method for binary classification. We use DWD to identify a linear combination of Φ-Space cell type scores such that cell-like objects from the two niches can be best separated. We implemented DWD using the kerndwd package [57, 58].
- *Partial least squares discriminant analysis (PLS-DA) for multiple niches* [22]. PLS-DA is a flexible method for multi-class classification. We use PLS-DA to identify linear combinations of Φ-Space cell type scores such that cell-like objects from multiple niches can be best separated. We implemented PLS-DA using the PhiSpace package [19].

Note that PLS-DA can also handle the case involving two niches. However, we found that the DWD loadings seemed to provide clearer biological interpretation in our case studies. Hence we opted for DWD for the two-niche case. We then rank the cell types according to their contribution towards predicting a given spatial niche, based on either the DWD or PLS-DA loadings. We define spatially enriched cell types as those that contribute most positively to predicting a specific niche.

#### Cell type co-presence (Fig 1C3)

To characterise the co-presence of different cell types at the same spatial location, we compute the co-presence matrix, defined as the correlation matrix of Φ-Space ST cell type scores. A positive correlation between two cell types indicates they tend to co-occur the same cell-like object. Tissue samples from donors with different disease conditions may exhibit very different cell type co-presence patterns. To facilitate a multi-sample analysis of these patterns, we apply PCA to reduce the dimensionality of cell type co-presence matrices, enabling visualisation and comparison across samples. We can also perform discriminant analysis using DWD or PLS-DA to identify cell type pairs whose co-presence distinguishes between disease conditions.

## Declarations

## Code availability

The Φ-Space R package is available on GitHub (https://github.com/jiadongm/PhiSpace), along with the R code for processing the data and reproduce our results.

## Data availability

The Human Lung Cell Atlas data (‘core’) [27] can be downloaded from CZ CELLxGENE (https://cellxgene.cziscience.com/collections/6f6d381a-7701-4781-935c-db10d30de293). The 10x Visium NSCLC dataset [24] is available at BioStudies (https://www.ebi.ac.uk/biostudies/) with accession number E-MTAB-13530.

The four scRNA-seq datasets from healthy and fibrotic lungs (immune, endothelial, epithelial and mesenchymal RDS objects) [28] are available at GEO with accession GSE227136. The CosMx NSCLC dataset (‘Processed Giotto Object’) [25] can be downloaded from https://nanostring.com/products/cosmx-spatial-molecular-imager/ffpe-dataset/nsclc-ffpe-dataset/.

The mouse spleen data from [29] is available at CNSA (CNGB Sequence Archive) of CNGBdb (https://db.cngb.org/cnsa/) under the accession number CNP0003930. The mouse bone marrow data from [30] is available at the Mouse HSC Atlas website (https://gillisweb.cshl.edu/HSC_atlas/). The mouse spleen CITE-seq data from [32] is available on GitHub (https://github.com/YosefLab/totalVI_reproducibility). The mouse neutrophil scRNA-seq data [31] is available at GEO under accession number GSE243466. The AML mouse spleen stereo-seq data [26] can be downloaded from Zenodo (https://zenodo.org/records/10685805) and the scRNA-seq data from the same spleen (‘Mouse 2’) [51] is available at GEO under accession number GSE161676.

## Acknowledgements.

We would like to thank Prof Mark Dawson, Dr Dane Vassiliadis and Ms Henrietta Holze (Sir Peter MacCullum Cancer Centre) for providing the processed AML mouse spleen Stereo-seq data and their helpful suggestions for writing the Stereo-seq case study. We would also like to thank Mr Yidi Deng (University of Melbourne) for helpful discussions.

