## Supplemental material for "Φ-Space ST: a platform-agnostic method to identify cell states in spatial transcriptomics studies"

### Supplementary materials for: Φ-Space ST: a sequencing platform-agnostic computational method for identifying cell states in spatial transcriptomics studies

Jiadong Mao[1], Jarny Choi[2, &], Kim-Anh Lê Cao[1,*,&]

[1]Melbourne Integrative Genomics, School of Mathematics and Statistics, The University of Melbourne, Australia

[2]Bioinformatics and Cellular Genomics, St Vincent's Institute, Australia

& indicates equal contribution

Throughout this document, unless otherwise indicated, we used the R package `PhiSpace` to produce case study results [1].

#### S1 Details of datasets and data preprocessing

**Human Lung Cell Atlas (HLCA).** We downloaded the 'Core' HLCA dataset [2] and used 'ann_finest_level' in the metadata as the cell type annotation. We then used `subsample` to subsample 10% of cells from each cell type. If 10% of a cell type contained fewer than 50 cells, we subsample 50 cells from that cell type. We did a log1p normalisation of the count matrix by `logTransf`. The feature importance scores were obtained using `tunePhiSpace`.

**10x Visium NSCLC data.** The Visium samples [3] were downloaded in h5 format and were read using `Seurat::Load10X_Spatial`, which was then converted to SingleCellExperiment (SCE) objects. All Visium samples were log1p normalised using `logTransf`.

**scRNA-seq data from healthy and fibrotic lungs.** We downloaded the four scRNA-seq datasets corresponding to cell lineages [4] and used 'manual_annotation_1' as the cell type labels. For each dataset, we subsampled 10% of each cell type using `subsample` with parameter `minCellNum = 50`. The raw counts were log1p normalised using `logTransf`. The feature importance scores were obtained using `tunePhiSpace`.

**CosMx NSCLC data.** The original `Giotto` [5] object was downloaded and converted to SCE objects. Each sample were log1p normalised using `logTransf`.

**Mouse spleen scRNA-seq data.** The scRNA-seq data from specific-pathogen-free (SPF) mice was downloaded [6]. The raw counts were scran normalised [7]. We used 'celltypes' in the metadata as the cell type annotation.

**Mouse bone marrow scRNA-seq data.** The mouse bone marrow data [8] contained 'cellType', which was used as the cell type annotation. We subsampled 10% of cells from each cell type using `subsample` with parameters `minCellNum = 50`. Since the raw counts were not present in the original dataset, we rank normalised the already normalised assay using `RankTransf`. Rank normalisation makes data normalised using different methods more comparable [9].

**Mouse spleen CITE-seq data.** The mouse CITE-seq data [10] contained cells from both mouse spleen and thymus. Only spleen cells were retained. There was a mild batch effects between the two experimental batches, which we removed using `batchelor::fastMNN` [11]. The reconstructed log normalised data were then used for annotation. We used 'cell_types' in the metadata as the cell type annotation.

**Mouse neutrophil scRNA-seq data.** The mouse neutrophil scRNA-seq data [12] contained neutrophils from different healthy and tumorous mouse tissues. The raw counts were scran normalised [7]. We used 'ManuscriptClusters' in the metadata as the cell type annotation.

**AML mouse spleen scRNA-seq data.** When using the mouse spleen [6], the mouse CITE-seq [10] and the neutrophil [12] datasets as references, we scran normalised [7] the AML mouse spleen scRNA-seq data [13]. When using the mouse bone marrow [8] as reference, we rank normalised the AML data to match the reference. When using the AML scRNA-seq data as reference to annotate the Stereo-seq data (see below), we log1p normalised the scRNA-seq data using `logTransf`.

**AML mosue spleen Stereo-seq data.** The bin50 Stereo-seq data [14] was log1p normalised using `logTransf`.

#### S2 Details of case studies

**Parameter tuning for Φ-Space.** In all the case studies, we used the default choice of `ncomp`, the number of PLS components, as in [15]. That is, we always let `ncomp` $= K$, with $K$ denoting the total number of cell types in the reference. In the CosMx case study, the query CosMx data only contained 960 genes, 951 of which were in the reference dataset. We did not do any gene selection and used all 951 shared genes for training Φ-Space. For the Visium and Stereo-seq case studies, we selected the top `nfeat` most useful features for predicting each cell type in Φ-Space ST, as in [15]. For the Visium case study, we selected `nfeat` $= 500$. In the Stereo-seq case study, 4 references were used and we selected `nfeat` $= 500$ for all references except Neutro; for Neutro we selected `nfeat` $= 600$. In both the Visium and Stereo-seq case studies, `nfeat` were so selected as to result in around $3,000$ genes for training Φ-Space.

**Implementation of deconvolution methods for Visium case study.** For each deconvolution method below, we used the same feature selection strategy as for Φ-Space ST above, to make sure that each deconvolution model was trained using the same number of features. We used the R package `spacexr` [16] to implement RCTD [17], which took unnormalised counts as input. We used the Python package `cell2location` (Version 0.1.4) to implement cell2location [18]. We followed the vignette (`https://cell2location.readthedocs.io/en/latest/notebooks/cell2location_tutorial.html`) for the training the the prediction, using tuning parameters specified in the vignette. We used 'q05_abundance' as the predicted cell type abundance. We used the Python package `tacco` (Version 0.2.2) to implement TACCO [19]. We used the `tacco.tl.annotate` function with default parameters.

**Number of meaningful markers in Visium case study.** In the Visium case study (Fig 2, for the prediction of cell type $C$ in tissue $T$ by method $M$, we defined the number of meaningful markers as the size of the intersection between the following two sets:

- *The spatial differentially expressed (DE) genes.* This gene set was defined as marker genes of the spatial region in tissue $T$ predicted to have higher $C$ identity by method $M$. We divided all Visium spots into two groups: the top 20% of spots having higher $C$ cell type scores, estimated by method $M$, and the remaining 80%. Then we used the R package MAST [20] to identify the DE genes in the first group of Visium spots. We ranked all DE genes according to their gene importance scores, as defined in [21]. We defined the top 100 genes as the spatial DE genes.

- *The cell type marker genes.* For downloaded the reference dataset, i.e. the human lung cell atlas [2], from the Azimuth project website [22], where 10 marker genes were used to define each cell type. To match the size of the spatial DE gene set above, we defined the marker genes of cell type $C$ as the 100 genes including the original 10 Azimuth markers plus the 90 genes that were most positively correlated with the Azimuth markers.

**Favourable prognosis scores of NSCLC samples.** In the Visium case study, for each cancerous sample, the favourable prognosis scores (see Fig 2) were calculated based on NSCLC prognosis markers identified in [23]. For LUSC, one favourable gene marker, CELSR2, was identified by [23]. The favourable prognosis score of a LUSC Visium sample was defined as the ratio between the total gene count of CELSR2 in that sample and the the mean total gene count averaged over all genes observed in that sample. For LUAD, 4 favourable markers (PERP, ELFN2, ARHGAP12 and QSOX1) and 6 unfavourable markers (KRT17, KRT6A, S100A2, TRIM29, REPS1 and GPC1) were identified by [23]. The favourable prognosis score of a LUAD Visium sample was defined as the ratio between the mean total gene count of the favourable markers and the mean total gene count of the unfavourable markers.

**Clustering Stereo-seq bins based on gene expression and Φ-Space score.** We clustered the Stereo-seq bins according to their gene expression profiles as follows. We computed the top 15 PCs of the log1p-transformed gene expression matrix and then applied $k$-means clustering to the 15 PCs by using the R function `kmeans` with the `algorithm="Lloyd"`, `iter.max=500` and `nstart=50`. We let the number of clusters $k = 8$, following [14]. We also computed the top 15 PCs of the Φ-Space scores and then applied $k$-means using the same specifications as above.

**Clustering Stereo-seq bins based on clonal composition.** To identify meta-clones, i.e. group of Stereo-seq bins with similar composition of individual AML clones, we designed a workflow as follows.

1. Generation of raw 'barcode-seq' data. We generated a sparse data matrix where columns were spatial bins and rows were individual AML barcodes. Each element of the matrix represented the count of a specific barcode detected in a specific bin.

2. Low abundance barcodes filtering. We filtered out barcodes with fewer than 50 total count.

3. Barcode-seq count matrix augmentation by local smoothing. That is, for each spatial bin, its neighbouring bins within the 500 µm diameter were identified and the average bin counts were computed as the augmented bin counts of that bin.

4. Meta-clone identification by clustering. We clustered the bins based on their augmented barcode counts by using $k$-means.

In the mouse spleen data analysed in the Stereo-seq case study, after the above filtering step, 35 out of 370 barcodes were retained. In the augmentation step, we varied the diameter values from 50 µm to 1000 µm and found that the clustering results were not sensitive to the diameter value, as long as it was not smaller than 100 µm. In the final clustering step, we varied the number of clusters $k$ from 2 to 10 and the clustering results were shown in Supplementary Fig S5. We presented the result for $k = 8$ in the main text (see Fig 4).

**Enriched cell types in AML meta-clones compared to background.** To find the enriched cell types in each AML meta-clones compared to the background (non-barcoded bins), we opt for a hypothesis-testing-based approach as follows. For each cell type, we compare the Φ-Space ST scores of bins in each meta-clone for that cell type to scores of bins in non-barcoded bins, by a two-sample Wilcoxon rank sum test using `stats::wilcox.test`. Then we calculated mean fold change (MFC) of a given cell type as

$$\text{MFC} = \frac{\text{mean cell type score of meta-clone} - \text{mean cell type score of background}}{\text{standard deviation of cell type score of background}}.$$

Finally, the significance score of a given cell type is calculated as

$$-\log(p) \times \text{MFC},$$

where $p$ denotes the p-value of the Wilcoxon test above. The sign of the significance score indicates if a cell type tends to be enriched (positive) or depleted (minus) in a meta-clone. For each, cell type, we selected five cell types that had the largest positive significance scores (all had adjusted p-value smaller than 0.05) as the meta-clone's most enriched cell types.

### S1 Supplementary figures

### S2 Supplementary tables

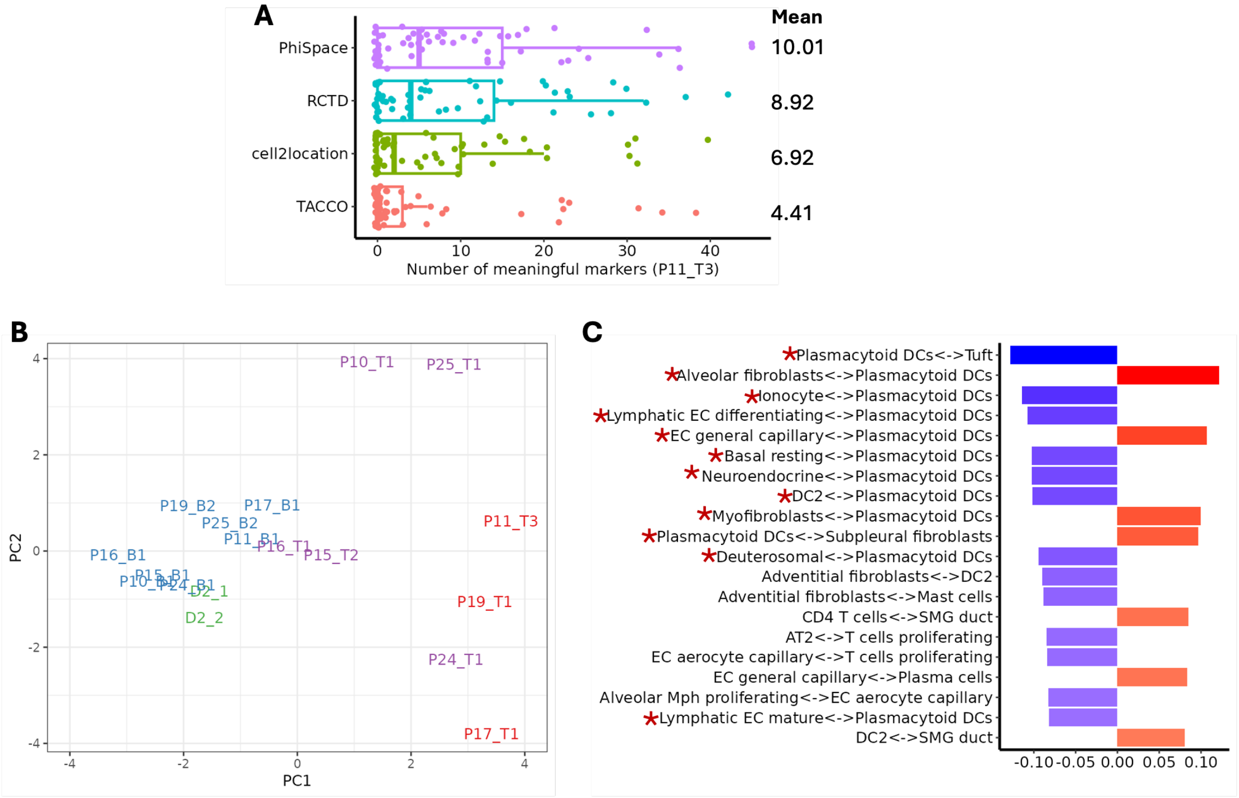


**Fig S1: Immune cell type interactions in 10x Visium lung cancer samples..** **A:** Boxplot comparing number of meaningful markers for the prediction of all cell types in P11_T3. **B:** Similar to Fig 2A, but now focusing on immune cell types, we filtered out correlations between non-immune cell types from the correlation matrices and recomputed their first two PCs. The separation between tumorous and non-tumorous tissues was largely retained compared to Fig 2A. **C:** Top 20 cell type interactions involving immune cell types contributing to PC1: 12/20 interactions (asterisks) involved plasmacytoid dendritic cells (pDCs).

[2] Lisa Sikkema, Ciro Ramírez-Suástegui, Daniel C Strobl, Tessa E Gillett, Luke Zappia, Elo Madissoon, Nikolay S Markov, Laure-Emmanuelle Zaragosi, Yuge Ji, Meshal Ansari, Marie-Jeanne Arguel, Leonie Apperloo, Martin Banchero, Christophe Bécavin, Marijn Berg, Evgeny Chichelnitskiy, Mei-I Chung, Antoine Collin, Aurore C A Gay, Janine Gote-Schniering, Baharak Hooshiar Kashani, Kemal Inecik, Manu Jain, Theodore S Kapellos, Tessa M Kole, Sylvie Leroy, Christoph H Mayr, Amanda J Oliver, Michael von Papen, Lance Peter, Chase J Taylor, Thomas Walzthoeni, Chuan Xu, Linh T Bui, Carlo De Donno, Leander Dony, Alen Faiz, Minzhe Guo, Austin J Gutierrez, Lukas Heumos, Ni Huang, Ignacio L Ibarra, Nathan D Jack-

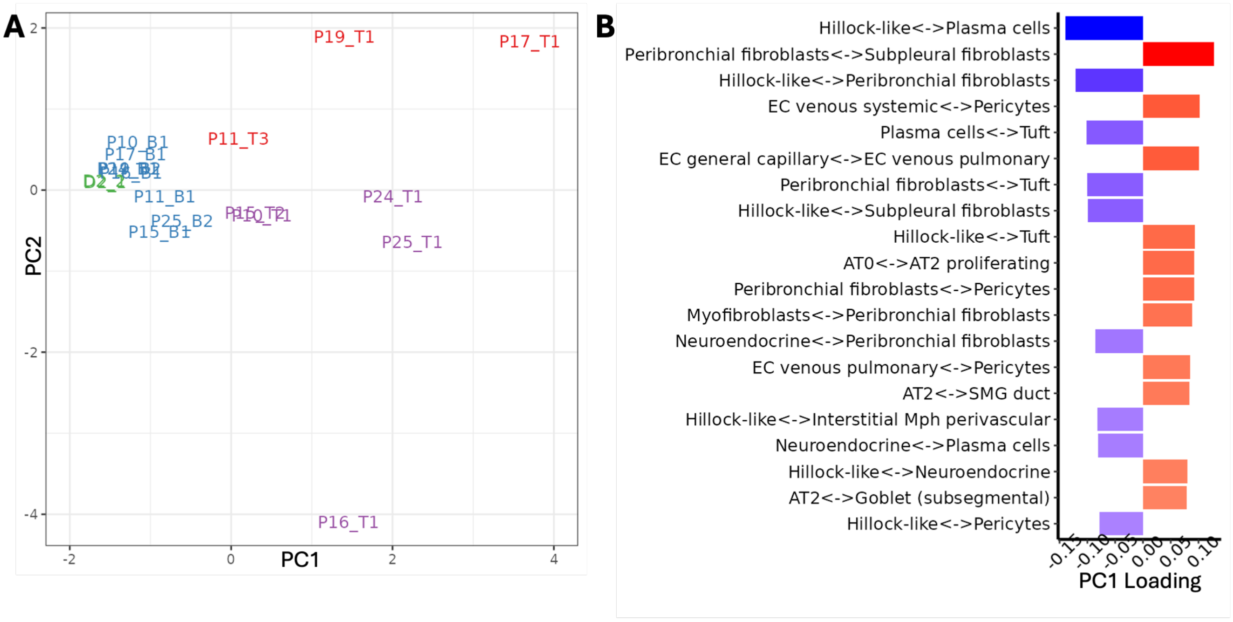


**Fig S2: Cell type co-presence analysis using RCTD deconvolution results.** **A:** The first two principal components (PCs) of cell type co-presence matrices derived using RCTD deconvolution results. We observed separation of different disease conditions. However, two PCs were needed for separating LUAD, LUSC and non-cancerous samples instead of just PC1 as in the Φ-Space ST results in Fig 2D. Moreover, neither of the RCTD PCs here seems to correspond to prognosis (Fig 2E). **B:** Top 20 loading values (absolute value) contributing to PC1. Fibroblasts still plays a prominant role, similar to the Φ-Space ST results as in Fig 2F.

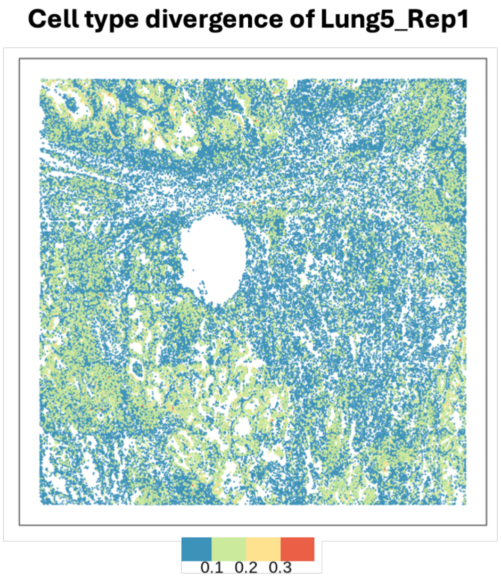


**Fig S3:** Cell type divergence of all segmented cells in Lung5_Rep1, computed based on their phenotype space embeddings derived using Φ-Space ST. The tumour and lymphoid structure domains in Fig 3A showed higher divergence.

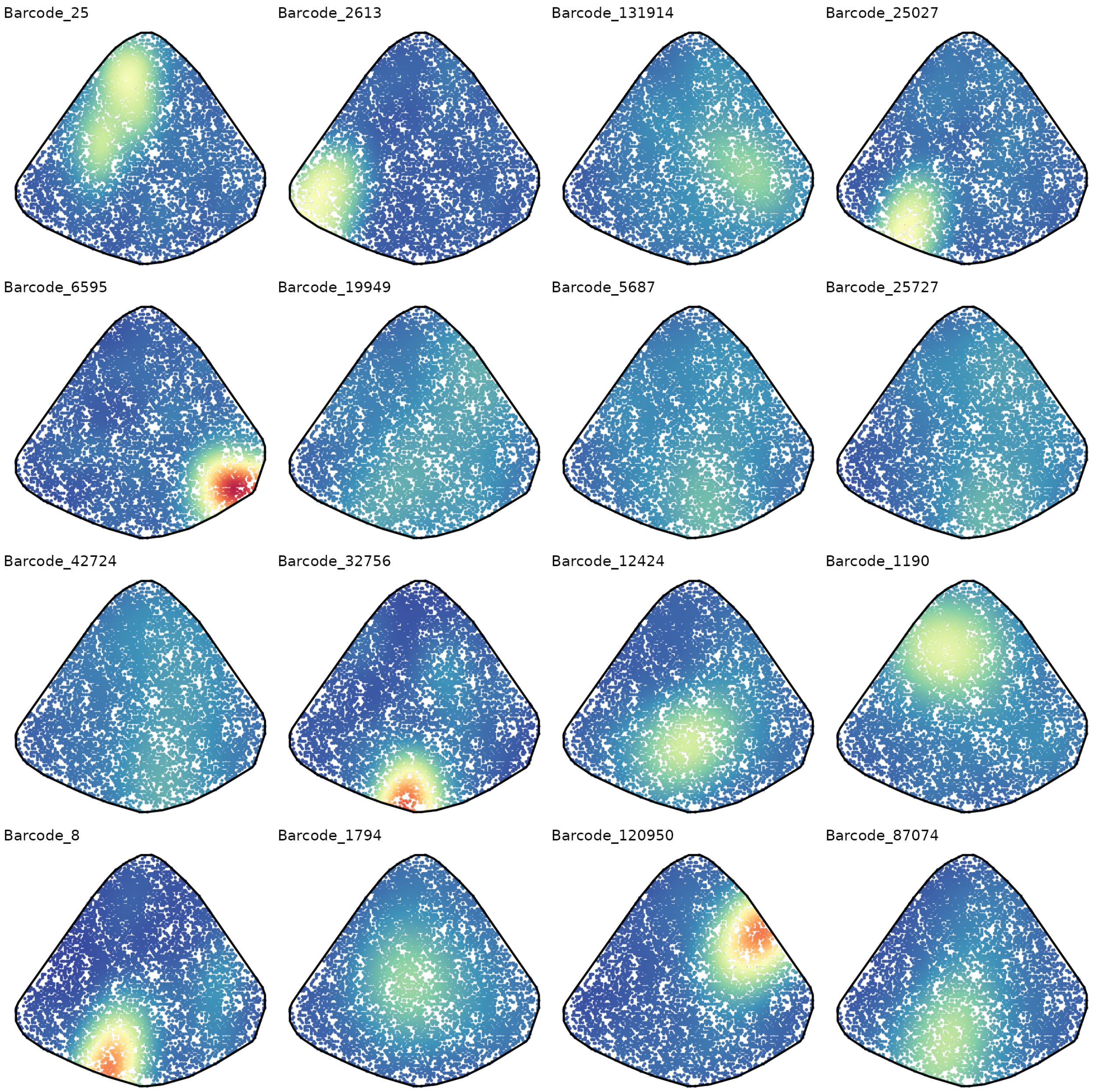


**Fig S4: Spatial distributions of most populous AML barcodes**. Each subfigure showed the spatial distribution of a specific AML barcode, where each spot represented a barcoded Stereo-seq bin. The spatial distribution of each barcode was estimated using a 2-dimensional kernel density estimator [24, 25]. The warmer the colour was, the higher the estimated density value.

son, Preetish Kadur Lakshminarasimha Murthy, Mohammad Lotfollahi, Tracy Tabib, Carlos Talavera-López, Kyle J Travaglini, Anna Wilbrey-Clark, Kaylee B Worlock, Masahiro Yoshida, Lung Biological Network Consortium, Maarten van den Berge, Yohan Bossé, Tushar J Desai, Oliver Eickelberg, Naftali Kaminski, Mark A Krasnow, Robert Lafyatis, Marko Z Nikolic, Joseph E Powell, Jayaraj Rajagopal, Mauricio Rojas, Orit Rozenblatt-Rosen, Max A Sei-

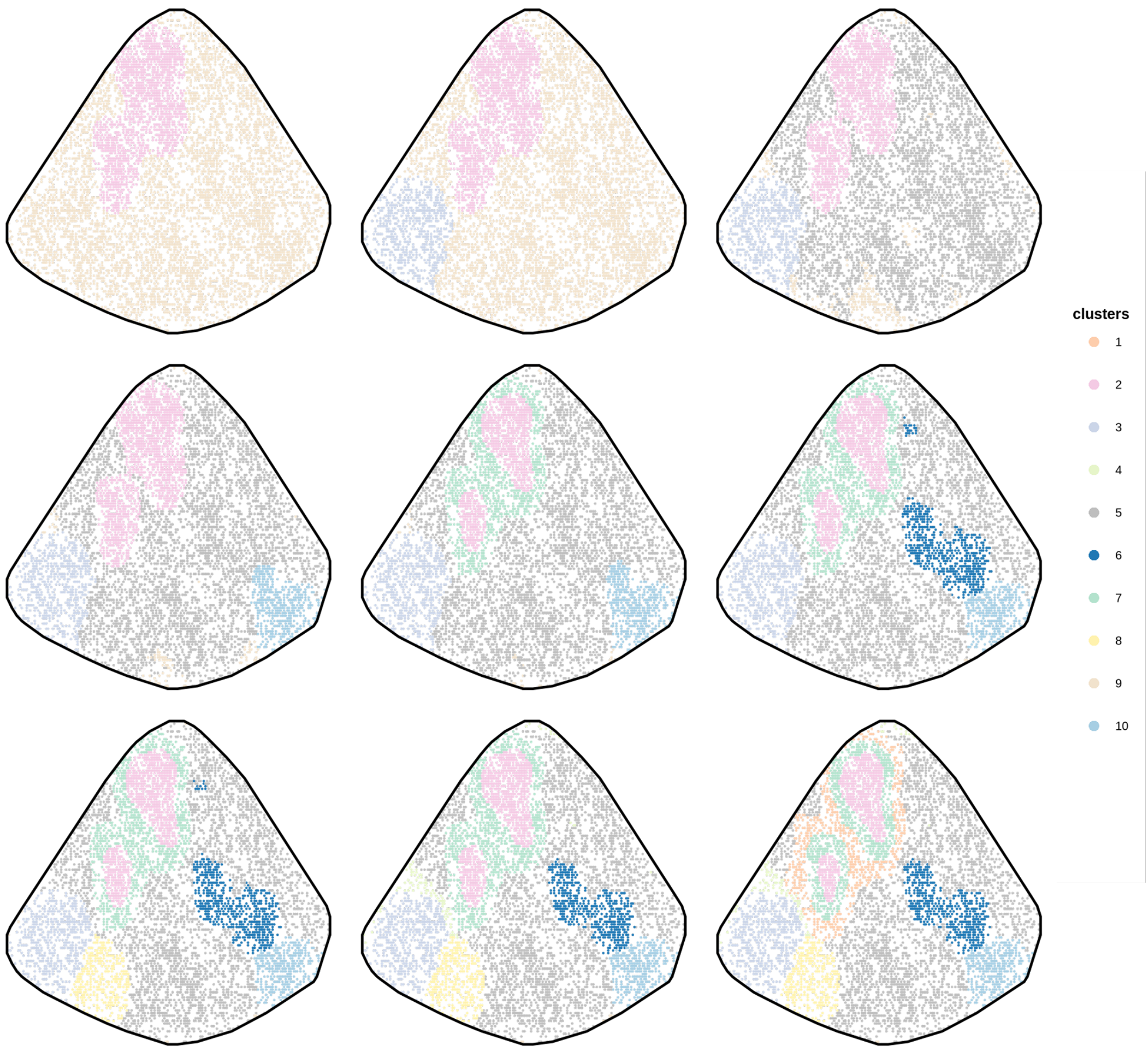


**Fig S5: Clustering Stereo-seq bins according to their barcode composition**. We varied the number of clusters $k$ from 2 to 10 in the meta-clone identification pipeline (see Supplementary Methods Section S2). Each subfigure above represented a specific choice of $k$.

bold, Dean Sheppard, Douglas P Shepherd, Don D Sin, Wim Timens, Alexander M Tsankov, Jeffrey Whitsett, Yan Xu, Nicholas E Banovich, Pascal Barbry, Thu Elizabeth Duong, Christine S Falk, Kerstin B Meyer, Jonathan A Kropski, Dana Pe'er, Herbert B Schiller, Purushothama Rao Tata, Joachim L Schultze, Sara A Teichmann, Alexander V Misharin, Martijn C Nawijn, Malte D Luecken, and Fabian J Theis. An integrated cell atlas of the lung in health and disease. *Nat. Med.*, 29(6):1563–1577, 2023.

[3] Marco De Zuani, Haoliang Xue, Jun Sung Park, Stefan C Dentro, Zaira Seferbekova, Julien

**Table S1: Niche-specific enriched cell types in all lung samples in CosMx case study.**

| | immune | lymphoid structure | macrophages | myeloid-enriched stroma | neutrophils | plasmablast-enriched stroma | stroma | tumor interior | tumor-stroma boundary |
|---|---|---|---|---|---|---|---|---|---|
| **Lung5_Rep1** | Alveolar Mph CCL3+ | B cells | Non-classical monocytes | Alveolar Mph CCL3+ | Alveolar Mph CCL3+ | Plasma cells | Adventitial fibroblasts | AT1 | AT1 |
| | B cells | CD4 T cells | Alveolar Mph CCL3+ | Non-classical monocytes | EC aerocyte capillary | Adventitial fibroblasts | Smooth muscle | Basal resting | Multiciliated (non-nasal) |
| | CD8 T cells | EC venous systemic | Monocyte-derived Mph | EC aerocyte capillary | Goblet (nasal) | Club (nasal) | Non-classical monocytes | Multiciliated (non-nasal) | Suprabasal |
| | EC aerocyte capillary | CD8 T cells | EC aerocyte capillary | NK cells | NK cells | Club (non-nasal) | DC2 | Suprabasal | Basal resting |
| | CD4 T cells | Hillock-like | NK cells | EC general capillary | EC general capillary | Smooth muscle | Alveolar fibroblasts | Hillock-like | Smooth muscle |
| **Lung5_Rep2** | CD4 T cells | B cells | | Alveolar Mph CCL3+ | Alveolar Mph CCL3+ | Plasma cells | Adventitial fibroblasts | AT1 | AT1 |
| | Suprabasal | CD4 T cells | | Non-classical monocytes | EC aerocyte capillary | Club (nasal) | Smooth muscle | Basal resting | Multiciliated (non-nasal) |
| | NK cells | EC venous systemic | | NK cells | NK cells | Adventitial fibroblasts | DC2 | Multiciliated (non-nasal) | Basal resting |
| | Alveolar Mph CCL3+ | CD8 T cells | | EC aerocyte capillary | Goblet (nasal) | EC general capillary | Lymphatic EC mature | Suprabasal | Suprabasal |
| | AT1 | EC venous pulmonary | | EC general capillary | AT1 | Alveolar fibroblasts | Alveolar macrophages | Hillock-like | Classical monocytes |
| **Lung5_Rep3** | CD4 T cells | B cells | Alveolar Mph CCL3+ | Alveolar Mph CCL3+ | Alveolar Mph CCL3+ | Plasma cells | Adventitial fibroblasts | AT1 | Smooth muscle |
| | Non-classical monocytes | CD4 T cells | EC aerocyte capillary | EC aerocyte capillary | EC aerocyte capillary | Club (nasal) | Smooth muscle | Basal resting | AT1 |
| | Alveolar Mph CCL3+ | EC venous systemic | NK cells | Non-classical monocytes | NK cells | AT2 | DC2 | Multiciliated (non-nasal) | Multiciliated (non-nasal) |
| | NK cells | pre-TB secretory | CD4 T cells | NK cells | Non-classical monocytes | Adventitial fibroblasts | Non-classical monocytes | Hillock-like | Basal resting |
| | EC aerocyte capillary | CD8 T cells | Goblet (nasal) | EC general capillary | EC general capillary | Plasmacytoid DCs | Alveolar fibroblasts | Suprabasal | Hillock-like |
| **Lung6** | Alveolar Mph CCL3+ | Alveolar Mph CCL3+ | Alveolar Mph CCL3+ | Alveolar Mph CCL3+ | | | Alveolar Mph CCL3+ | Hillock-like | CD4 T cells |
| | Interstitial Mph perivascular | Plasma cells | Interstitial Mph perivascular | Interstitial Mph perivascular | | | Plasma cells | NK cells | Alveolar Mph CCL3+ |
| | Plasma cells | Interstitial Mph perivascular | Alveolar macrophages | DC2 | | | Interstitial Mph perivascular | Suprabasal | Classical monocytes |
| | DC2 | CD4 T cells | DC2 | AT2 | | | CD4 T cells | Basal resting | Lymphatic EC mature |
| | CD4 T cells | B cells | Monocyte-derived Mph | Plasma cells | | | Lymphatic EC mature | Goblet (nasal) | Plasma cells |
| **Lung9_Rep1** | EC venous pulmonary | CD4 T cells | Monocyte-derived Mph | EC venous pulmonary | Monocyte-derived Mph | Plasma cells | EC venous pulmonary | pre-TB secretory | Adventitial fibroblasts |
| | Plasma cells | Plasma cells | Non-classical monocytes | EC arterial | EC venous pulmonary | B cells | EC aerocyte capillary | Neuroendocrine | Alveolar fibroblasts |
| | CD4 T cells | EC venous pulmonary | EC venous pulmonary | Non-classical monocytes | CD4 T cells | Smooth muscle | SMG duct | Deuterosomal | Smooth muscle |
| | EC aerocyte capillary | EC aerocyte capillary | EC arterial | SMG duct | Basal resting | EC venous pulmonary | EC general capillary | Tuft | Club (non-nasal) |
| | Alveolar fibroblasts | Lymphatic EC mature | SMG duct | EC aerocyte capillary | AT2 | Multiciliated (non-nasal) | Alveolar Mph CCL3+ | Hillock-like | EC general capillary |
| **Lung9_Rep2** | Alveolar fibroblasts | Alveolar fibroblasts | | Alveolar fibroblasts | Non-classical monocytes | Plasma cells | Alveolar fibroblasts | pre-TB secretory | Alveolar fibroblasts |
| | Plasma cells | Plasma cells | | DC2 | DC2 | Alveolar fibroblasts | Adventitial fibroblasts | Neuroendocrine | Plasma cells |
| | Non-classical monocytes | Non-classical monocytes | | Non-classical monocytes | Alveolar fibroblasts | Non-classical monocytes | DC2 | Hillock-like | Adventitial fibroblasts |
| | Adventitial fibroblasts | Adventitial fibroblasts | | Adventitial fibroblasts | Alveolar Mph CCL3+ | Adventitial fibroblasts | Smooth muscle | Deuterosomal | EC general capillary |
| | EC general capillary | DC2 | | Multiciliated (nasal) | Multiciliated (nasal) | Smooth muscle | Alveolar Mph CCL3+ | Tuft | Alveolar macrophages |
| **Lung12** | Monocyte-derived Mph | Plasma cells | | Plasma cells | CD4 T cells | Plasma cells | Adventitial fibroblasts | Goblet (nasal) | AT1 |
| | Suprabasal | B cells | | CD4 T cells | NK cells | B cells | Alveolar fibroblasts | AT1 | Club (nasal) |
| | EC arterial | Monocyte-derived Mph | | NK cells | CD8 T cells | EC venous pulmonary | EC general capillary | Classical monocytes | AT2 |
| | NK cells | EC venous pulmonary | | Non-classical monocytes | Non-classical monocytes | EC arterial | Alveolar macrophages | Monocyte-derived Mph | Classical monocytes |
| | CD8 T cells | EC arterial | | EC venous systemic | DC2 | Lymphatic EC mature | Smooth muscle | Basal resting | Smooth muscle |
| **Lung13** | Interstitial Mph perivascular | | Monocyte-derived Mph | Monocyte-derived Mph | Club (nasal) | | Adventitial fibroblasts | pre-TB secretory | AT2 |
| | Plasma cells | | Non-classical monocytes | pre-TB secretory | pre-TB secretory | | Suprabasal | Club (nasal) | Goblet (nasal) |
| | DC2 | | Club (non-nasal) | Suprabasal | Multiciliated (non-nasal) | | EC general capillary | Goblet (nasal) | AT1 |
| | EC general capillary | | Alveolar fibroblasts | CD8 T cells | CD4 T cells | | Alveolar fibroblasts | EC arterial | Classical monocytes |
| | CD8 T cells | | Pericytes | EC arterial | Suprabasal | | Interstitial Mph perivascular | AT1 | Club (nasal) |


Tessier, Sandra Curras-Alonso, Angela Hadjipanayis, Emmanouil I Athanasiadis, Moritz Gerstung, Omer Bayraktar, and Ana Cvejic. Single-cell and spatial transcriptomics analysis of non-small cell lung cancer. *Nat. Commun.*, 15(1):4388, 2024.

[4] Heini M Natri, Christina B Del Azodi, Lance Peter, Chase J Taylor, Sagrika Chugh, Robert Kendle, Mei-I Chung, David K Flaherty, Brittany K Matlock, Carla L Calvi, Timothy S Blackwell, Lorraine B Ware, Matthew Bacchetta, Rajat Walia, Ciara M Shaver, Jonathan A Kropski, Davis J McCarthy, and Nicholas E Banovich. Cell-type-specific and disease-associated expression quantitative trait loci in the human lung. *Nat. Genet.*, 56(4):595–604, 2024.

[5] Ruben Dries, Qian Zhu, Rui Dong, Chee-Huat Linus Eng, Huipeng Li, Kan Liu, Yuntian Fu, Tianxiao Zhao, Arpan Sarkar, Rani E George, Nico Pierson, Long Cai, and Guo-Cheng Yuan. Giotto, a toolbox for integrative analysis and visualization of spatial expression data. *bioRxiv*, 2020. doi: 10.1101/701680. URL `https://doi.org/10.1101/701680`.

[6] Yin Zhang, Juan Shen, Wei Cheng, Bhaskar Roy, Ruizhen Zhao, Tailiang Chai, Yifei Sheng, Zhao Zhang, Xueting Chen, Weiming Liang, Weining Hu, Qijun Liao, Shanshan Pan, Wen Zhuang, Yangrui Zhang, Rouxi Chen, Junpu Mei, Hong Wei, and Xiaodong Fang. Microbiota-mediated shaping of mouse spleen structure and immune function characterized by scRNA-seq and stereo-seq. *J. Genet. Genomics*, 50(9):688–701, 2023.

[7] Aaron T L Lun, Karsten Bach, and John C Marioni. Pooling across cells to normalize single-cell RNA sequencing data with many zero counts. *Genome Biol.*, 17:75, 2016.

[8] Benjamin D Harris, John Lee, and Jesse Gillis. A meta-analytic single-cell atlas of mouse bone marrow hematopoietic development. *bioRxiv*, page 2021.08.12.456098, 2021.

[9] Yanxiang Deng, Marek Bartosovic, Sai Ma, Di Zhang, Petra Kukanja, Yang Xiao, Graham Su, Yang Liu, Xiaoyu Qin, Gorazd B Rosoklija, Andrew J Dwork, J John Mann, Mina L Xu, Stephanie Halene, Joseph E Craft, Kam W Leong, Maura Boldrini, Gonçalo Castelo-Branco, and Rong Fan. Spatial profiling of chromatin accessibility in mouse and human tissues. *Nature*, 609(7926):375–383, 2022.

[10] Adam Gayoso, Zoë Steier, Romain Lopez, Jeffrey Regier, Kristopher L Nazor, Aaron Streets, and Nir Yosef. Joint probabilistic modeling of single-cell multi-omic data with totalVI. *Nat. Methods*, 18(3):272–282, 2021.

[11] Laleh Haghverdi, Aaron T. L. Lun, Michael D. Morgan, and John C. Marioni. Batch effects in single-cell RNA-sequencing data are corrected by matching mutual nearest neighbors. *Nat. Biotechnol.*, 36(5):421–427, 2018. doi: 10.1038/nbt.4091.

[12] Melissa S F Ng, Immanuel Kwok, Leonard Tan, Changming Shi, Daniela Cerezo-Wallis, Yingrou Tan, Keith Leong, Gabriel F Calvo, Katharine Yang, Yuning Zhang, Jingsi Jin, Ka Hang Liong, Dandan Wu, Rui He, Dehua Liu, Ye Chean Teh, Camille Bleriot, Nicoletta Caronni, Zhaoyuan Liu, Kaibo Duan, Vipin Narang, Iván Ballesteros, Federica Moalli, Mengwei Li, Jinmiao Chen, Yao Liu, Lianxin Liu, Jingjing Qi, Yingbin Liu, Lingxi Jiang, Baiyong Shen, Hui Cheng, Tao Cheng, Veronique Angeli, Ankur Sharma, Yuin-Han Loh, Hong Liang Tey, Shu Zhen Chong, Matteo Iannacone, Renato Ostuni, Andrés Hidalgo, Florent Ginhoux, and Lai Guan Ng. Deterministic reprogramming of neutrophils within tumors. *Science*, 383(6679): eadf6493, 2024.

[13] Katie A Fennell, Dane Vassiliadis, Enid Y N Lam, Luciano G Martelotto, Jesse J Balic, Sebastian Hollizeck, Tom S Weber, Timothy Semple, Qing Wang, Denise C Miles, Laura MacPherson, Yih-Chih Chan, Andrew A Guirguis, Lev M Kats, Emily S Wong, Sarah-Jane Dawson, Shalin H Naik, and Mark A Dawson. Non-genetic determinants of malignant clonal fitness at single-cell resolution. *Nature*, 601(7891):125–131, 2022.

[14] Henrietta Holze, Laure Talarmain, Katie A Fennell, Enid Y Lam, Mark A Dawson, and Dane Vassiliadis. Analysis of synthetic cellular barcodes in the genome and transcriptome with BARtab and bartools. *Cell Rep Methods*, page 100763, 2024.

[15] Jiadong Mao, Yidi Deng, and Kim-Anh Lê Cao. $\phi$-space: Continuous phenotyping of single-cell multi-omics data. *bioRxiv*, page 2024.06.19.599787, 2024.

[24] José E Chacón and Tarn Duong. *Multivariate kernel smoothing and its applications.* CRC Press, London, England, 1st edition edition, 2020.

[25] Tarn Duong. *ks: Kernel Smoothing*, 2024. URL `https://CRAN.R-project.org/package=ks`. R package version 1.14.2.
